# The Subsystem Mechanism of Default Mode Network Underlying Rumination: a Reproducible Neuroimaging Study

**DOI:** 10.1101/841239

**Authors:** Xiao Chen, Ning-Xuan Chen, Yang-Qian Shen, Hui-Xian Li, Le Li, Bin Lu, Zhi-Chen Zhu, Zhen Fan, Chao-Gan Yan

## Abstract

Rumination is a repetitive self-referential thinking style and posited to be an expression of abnormalities in the default mode network (DMN) in major depressive disorder (MDD). Recent evidences indicate DMN is not a unitary network but can be further divided into 3 functionally heterogenous subsystems. However, the subsystem mechanism through which DMN underlie rumination remain unclear. Here, with a modified continuous state-like paradigm, we induced healthy participants to ruminate or imagine objective scenarios (as a distraction control condition) on 3 different MRI scanners. We compared functional connectivities (FC) and inter-subject correlations (ISC) of the DMN and its 3 subsystems between rumination and distraction states. Results yielded a highly reproducible and dissociated pattern. During rumination, within-DMN FC was generally decreased compared to the distraction state. At the subsystem level, we found increased FC between the core and medial temporal lobe (MTL) subsystem and decreased FC between the core and dorsal medial prefrontal cortex (DMPFC) subsystem and within the MTL subsystem. Furthermore, we found decreased ISC within the MTL subsystem. These results suggest a specific and less synchronous activity pattern of DMN during rumination and shed new light on the association between rumination and DMN substrates regarding MDD.

## Introduction

Rumination is generally defined as a recurrent and passive focus on depressed mood itself and its possible causes and consequences (Nolen-Hoeksema S and J Morrow 1991). As a specific form of self-generated thoughts, rumination is common when people are expressing distress and is characterized by self-focused, past-oriented, negative thinking forms, highly automatic dynamic phenomenology, and contents of thought restricted to narrow themes (Watkins E et al. 2005; Fox KCR et al. 2018). Studies have implicated its pivotal role in the onset, maintenance as well as phenomenology of major depressive disorder (MDD) (Spasojević J and LB Alloy 2001; Nolen-Hoeksema S et al. 2008). Moreover, rumination is also deemed as the psychological expression of MDD patients’ alterations within the default mode network (DMN) (Kaiser RH et al. 2015). Indeed, convergent lines of evidence suggest the DMN may be the key brain network supporting rumination (Zhou HX et al. 2019). Using typical task-based designs, researchers found DMN region activities were higher in rumination compared to distraction, which is a control condition requiring participants to focus on externally oriented thoughts that are unrelated to themselves (Nolen-hoeksema S and J Morrow 1993; Johnson MK et al. 2006; Johnson MK et al. 2009; Cooney RE et al. 2010; Burkhouse KL et al. 2016). Another line of studies (Greicius MD et al. 2007; Zhu X et al. 2012; Lois G and M Wessa 2016; Zhu X et al. 2017) revealed that the strength of functional connectivity (FC) within the DMN is associated with subjects’ rumination traits in the resting state.

The DMN is a set of brain regions showing reduced activity during active attention demanding tasks (Buckner RL and DC Carroll 2007; Andrews-Hanna JR, JS Reidler, J Sepulcre, et al. 2010; Buckner RL and LM DiNicola 2019). Systematic reviews and meta-analyses implicated the DMN in a number of different processes including remembering the past, scene construction and theory of mind (Buckner RL and DC Carroll 2007; Spreng RN et al. 2009). These processes are posited to be forms of self-generated thought, with mental contents dependent on internally constructed rather than externally constructed representations (Christoff K et al. 2016; Buckner RL and LM DiNicola 2019). Further evidence suggests these processes may correspond to 3 heterogeneous subsystems: the core DMN subsystem, the dorsal medial prefrontal cortex (DMPFC) subsystem and the medial temporal lobe (MTL) subsystem (Andrews-Hanna JR 2012). In general, the core subsystem is proposed to be associated with self-referential processes and acts as the hub linking all three subsystems. The DMPFC subsystem mainly participates in theory of mind and mental simulations whereas the MTL subsystem plays an important role in autobiographical memory and generating novel thinking ensembles (Andrews-Hanna JR, JS Reidler, C Huang, et al. 2010; Andrews-Hanna JR 2012; Andrews-Hanna JR et al. 2014). These DMN brain regions are proposed to work collectively and adaptively to facilitate different forms of self-generated thoughts (Buckner RL and DC Carroll 2007; Buckner RL and LM DiNicola 2019). As a distinct form of self-generated thoughts, rumination has been proposed to be driven by specific interactions among subsystems of the DMN (Bar M 2009; Christoff K *et al*. 2016). However, such proposals still lack direct empirical evidence.

Although resting-state fMRI studies have been utilized to study the rumination trait across participants (Kaiser RH et al. 2016), rumination task fMRI studies are more suitable to reveal DMN patterns during active rumination states. Due to its repetitive and self-perpetuating nature (Lyubomirsky S and S Nolen-Hoeksema 1993), rumination is particularly suitable for a continuous semi-task paradigm, in which participants are asked to maintain a certain kind of mental process for a relatively long period (Milazzo AC et al. 2016). Several studies have used this kind of paradigm to explore the network mechanisms of rumination. Berman et al. (Berman MG et al. 2014) induced MDD patients and healthy controls to memorize certain negative autobiographical events continuously for 6 minutes as a rumination induction. They compared FC between posterior cingulate cortex (PCC) and the rest of the whole brain in rumination and in typical resting state, and found enhanced FC strength during rumination compared to resting-state across MDD patients and healthy controls. In another study (Zamoscik V et al. 2014), MDD patients and healthy controls were induced either to ruminate or be distracted for 4.5 minutes after a negative autobiographical recall, but no FC difference between rumination and distraction was found. With support vector machine, Milazzo et al. (Milazzo AC *et al*. 2016) investigated the differences in FC strengths between rumination and other mental states and found FC involving amygdala best distinguished rumination. Intriguing but inconsistent, these studies have demonstrated the feasibility of the rumination state paradigm, but the heterogeneity in rumination induction strategies and control conditions suggests a more rigorous approach is required to investigate the DMN mechanisms underlying rumination.

Apart from FC, inter-subject correlations (ISC) have been found to be useful and reliable in detecting stimulus-driven responses within the DMN (Hasson U et al. 2004; Hasson U et al. 2010). A recent study found ISC may also represent shared internal mental processes (Chen J et al. 2017). Rumination is characterized by its specific phenomenological features: diverse in contents but shared in forms (Watkins E *et al*. 2005; Andrews-Hanna J et al. 2013), which could be reflected by ISC alterations. Another concern in the field is the lack of reproducibility of small sample size, hypothesis-driven fMRI studies (Button KS et al. 2013; Poldrack RA et al. 2017). As one recent validation research showed, different researchers tend to report substantially variable binary results, even when they are analyzing identical datasets and testing same hypothesis(Botvinik-Nezer R et al. 2019). Whereas practices such as data sharing, transparent study reporting and enlarging sample size also help (Poldrack RA *et al*. 2017), we addressed this issue straightforwardly by repetitively scanning all participants on 3 different scanners in counterbalanced order and report only reproducible results. Besides, we openly shared all raw neuroimaging data and scripts so that our findings can be validated by other interested researchers. We hope these practices can further enhance the reproducibility of our results.

The present study sought to directly investigate the interaction pattern (FCs) and shared neural process (ISCs) within DMN when subjects are actively engaging in rumination. To do so, we 1) designed a modified continuous, self-driven rumination state task that can provide state-like data suitable for FC and ISC analysis; 2) delineated the DMN into 3 subsystems of core, DMPFC, MTL according to Andrews-Hanna and colleagues’ studies (Andrews-Hanna JR, JS Reidler, J Sepulcre, *et al*. 2010; Andrews-Hanna JR *et al*. 2014) and analyzed FCs and ISCs at the subsystem level; 3) assured reproducibility by scanning participants on three distinct scanners in counterbalanced order. We hypothesized that FC between core and MTL subsystems would be enhanced and ISCs within DMN subsystems would be altered during rumination.

## Materials and Methods

### Participants

We recruited 42 young healthy adult participants from the community. One participant dropped out, leaving a final sample of 41 (22 females; mean age = 22.7 ± 4.1 years; 40 right-handed and one left-handed) subjects. The current study was approved by the Institutional Review Board of the Institute of Psychology, Chinese Academy of Sciences. All participants gave informed consent.

### Experimental procedure

A few days before the first scan, participants were invited to take part in an interview. During the interview, they first generated key words for individualized negative autobiographical events, then pre-experienced the rumination induction task. They also filled out a series of scales.

Several days before the first scan, participants were invited to take part in a brain storm outside scanner. Prior to this session, we briefly introduced the purpose and the mental states we would like them to engage. Rumination was stated as a “passive and repetitive thinking on the negative events themselves and their possible consequences” while distraction was introduced as an “imagery imagination that is unrelated to ourselves”. Subjects were required to try to stay in each mental state and keep thinking or imaging as the stimuli were displayed on the screen. We told them “we were interested in people’s coping strategies when they encounter negative life events and the brain mechanisms supporting them”. Participants then were prompted to briefly recall 8 negative autobiographical events (no time limits, but most participants spent about 10-15s on each event) in their life. After that, participants rated each event with a series of 9-point Likert scales like “How easily you recall that event?” or “Can you recall that event vividly now?” According to participants’ ratings, 4 highest scored events would be picked. Then participants were asked to recall each event as vividly as possible for 2 minutes. This was also a practice for them. After recalling, participants had to generate 3 to 5 key words for each event. These key words will be presented in the fMRI task to help them recall the negative events, and thus induce negative affect. Besides, participants also pre-experienced the rumination and distraction states.

Participants completed identical fMRI task on 3 distinct MRI scanners. All fMRI tasks include 4 different sessions: resting state, negative events recall, rumination state and distraction state. An 8-minute resting state came first as a baseline. In this session, participants were told to look at a fixation cross on the screen and not to think anything in particular. Then participants would recall negative autobiographical events which was prompted by a series of keywords generated by themselves during the “brain storm” stage on the screen. Participants were told to recall as vividly as they could and imagine they were experiencing those negative events again. Negative autobiographical events recall was also used as an affect induction to help participants enter a dysphoric mood condition so that they could be induced into rumination more easily. After negative autobiographical recall, participants sequentially went through rumination state and distraction state whose order was counter-balanced across subjects. In the rumination state, questions such as “Think: Analyze your personality to understand why you feel so depressed in the events you just memorized” were presented on the screen to help participants think about themselves, while in distraction state, prompts like “think: The layout of a typical classroom” were presented to help participants focus on an objective and concrete scenery (Table S1). These items were obtained and modified from the Rumination Induction Task (RIT) (Lyubomirsky S and S Nolen-Hoeksema 1995) and RRS which were widely used and proved to be effective to induce participants into rumination and distraction states (Lyubomirsky S and S Nolen-Hoeksema 1993; Nolen-hoeksema S and J Morrow 1993; Huffziger S and C Kuehner 2009). All mental states (negative events recall, rumination and distraction) except for the resting state contained four randomly sequentially presented stimuli (keywords or mental prompts). Each one lasted for 2 minutes then switched to next without any inter stimuli intervals (ISI), forming an 8-minute continuous mental states (see Figure S1). Our pre-study showed that subject would easily keep on thinking with one single prompt for about 2 minutes, so we divided each mental states into 4 blocks to help participants stay in the mental state (see Figure S2). Of note, before resting state and after each mental state, participants needed to report their present emotion state (happy vs. sad) with a 9-point Likert scale. Apart from this, after finishing each mental state, they also needed to report the contents they just thought with a self-report questionnaire. The items of this questionnaire were derived from a factor analysis regarding the contents and form of individuals self-generated thoughts (see Table S2) (Gorgolewski KJ et al. 2014).

### Data acquisition

The scan parameters used in the present research were the recommended standardized sequence of the Association of Brain Imaging (http://www.abimaging.org). These parameters were developed to harmonious site effects from different models of scanners. Images were acquired on two 3 Tesla GE MR750 scanners at Magnetic Resonance Imaging Research Center, Institute of Psychology, Chinese Academy of Sciences (henceforth IPCAS) and Peking University with 8-channel head-coils. Another 3 Tesla SMENENS PRISMA scanner (henceforth PKUSIEMENS) with an 8-channel head-coil in Peking University was also used for image acquisition. The tasks were identical on different scanners and the order of three scans was counter-balanced across participants. Before functional images acquisitions, all participants underwent a 3D T1-weighted scan first (IPCAS/PKUGE: 192 sagittal slices, TR = 6.7 ms, TE = 2.90 ms, slice thickness/gap = 1/0mm, in-plane resolution = 256 × 256, inversion time (IT) = 450ms, FOV = 256 × 256 mm, flip angle = 7°, average = 1; PKUSIEMENS: 192 sagittal slices, TR = 2530 ms, TE = 2.98 ms, slice thickness/gap = 1/0 mm, in-plane resolution = 256 × 224, inversion time (TI) = 1100 ms, FOV = 256 × 224 mm, flip angle = 7°, average=1). After T1 image acquisition, functional images were scanned for all three mental states (IPCAS/PKUGE: 33 axial slices, TR = 2000 ms, TE = 30 ms, FA = 90°, thickness/gap = 3.5/0.6 mm, FOV = 220 × 220 mm, matrix = 64 × 64; PKUSIEMENS: 62 sagittal slices, TR = 2000 ms, TE = 30 ms, FA = 90°, thickness = 2 mm, Slice acceleration = 2, FOV = 224 × 224 mm). Multiband scan (slice acceleration = 2) was performed on PKUSIEMENS to get a finer voxel size of 2 × 2 × 2 mm.

### MRI data analysis

Data preprocessing was done with Data Processing Assistant for Resting-State fMRI (DPARSF) (Yan C-G and Y-F Zang 2010). We first excluded any participant with > 0.2mm average frame displacement (FD) calculated following Jenkinson et al. (Jenkinson M et al. 2002; Parkes L et al. 2018), but no participant was removed. Then the initial 5 volumes were discarded, and slice-timing correction was performed. The time series of images for each subject were realigned using a six-parameter (rigid body) linear transformation with a two-pass procedure (registered to the first image and then registered to the mean of the images after the first realignment). After realignment, individual T1-weighted MPRAGE images were co-registered to the mean functional image using a 6 degree-of-freedom linear transformation without re-sampling and then segmented into gray matter, white matter (WM), and cerebrospinal fluid (CSF). Finally, transformations from individual native space to MNI space were computed with the Diffeomorphic Anatomical Registration Through Exponentiated Lie algebra (DARTEL) tool(Ashburner J 2007).

To minimize head motion confounds, we utilized the Friston 24-parameter model (Friston KJ et al. 1996) to regress out head motion effects. The Friston 24-parameter model (i.e., 6 head motion parameters, 6 head motion parameters one time point before, and the 12 corresponding squared items) was chosen based on prior work that higher-order models remove head motion effects better (Satterthwaite TD et al. 2013; Yan C-G et al. 2013). Additionally, mean FD was used to address the residual effects of motion in group analyses. Mean FD is derived from Jenkinson’s relative root mean square (RMS) algorithm (Jenkinson M *et al*. 2002). As global signal regression (GSR) is still a controversial practice in the R-fMRI field(Murphy K and MD Fox 2016), we did not perform GSR. Other sources of spurious variance (WM and CSF signals) were also removed from the data through linear regression to reduce respiratory and cardiac effects. Additionally, linear trends were included as a regressor to account for drifts in the blood oxygen level dependent (BOLD) signal. For mental states (rumination and distraction), additional regressors representing four sets of stimuli were further included to regress out the impacts of the switch of stimuli. We performed temporal bandpass filtering (0.01-0.1Hz) on all time series. Lastly, all functional images were smoothed with a 6 mm FWHM Gaussian kernel.

We defined a total of 24 anatomical regions of interest (ROIs), which is based on Yeo’s 17-network parcellation derived from the data of 1000 young healthy participants (Yeo BT et al. 2011). These 24 ROIs can be further divided into 3 subsystems (Table S3). For all ROIs, spatially connected regions were combined to form a single ROI, whereas spatially disconnected regions became separated ROIs. The original atlas was converted to the MNI space. We firstly extracted the average time series from each ROI and calculated FC of each ROI-pair, forming a 24 × 24 FC matrix. We then Z-transformed all FCs with a Fisher’s r-to-z transformation. For DMN FC, we simply average across all FCs in the matrix. For FC within each subsystem, we average FCs among ROIs of the same subsystem and all the FCs reflecting pair-wise connections between subsystems as FCs between subsystems. These values and FCs of all ROI pairs were then submitted to a general linear model to conduct a two-tailed paired t test, with head motion of each states regressed out (n = 41). We used Bonferroni correction to control the subsystem-level multiple comparisons and FDR correction for ROI-level multiple comparisons.

We calculated ISC on the ROI level. Because all mental states contain four 2-minute blocks, we split all states’ time series of each ROI into four segments according to the stimuli. We then separately calculated ISC for each segment by correlating each subjects’ time course to the average of others (N – 1) in the same ROI. The initial 5 time points were discarded for the first stimuli, so we calculated the correlation between the remain 55 time points for the first stimuli with the latter 55 time points of the average time courses. After Fisher’s r-to-z transformation, we average all ISCs across DMN and all 3 subsystems as the system level ISC. Two-tailed paired t tests were conducted to compare ISCs of rumination and distraction states (n = 41).

### Data availability

All neuroimaging data will be shared through the R-fMRI Maps project (http://rfmri.org/maps) upon publication. Analysis codes are made available online (https://github.com/Chaogan-Yan/PaperScripts/tree/master/Chen_2019).

## Results

### Behavioral Results

Behavioral results revealed an extraordinarily reproducible pattern (Figure 1). Compared to distraction and resting states, the contents of rumination were more about past, self, others, in speech form, and made participants sadder (p < 0.05, Bonferroni corrected, Table S4). After negative autobiographical memory induction, participants’ emotional ratings were significantly decreased (IPCAS: t = −6.32, p < 0.001, df = 40; PKUGE: t = −7.71, p < 0.001, df = 40; PKUSIEMENS: t = −6.28, p < 0.001, df = 40) compared to baseline and remained decreased after rumination (IPCAS: t = −4.44, p < 0.001, df = 40; PKUGE: t = −4.39, p < 0.001, df = 40; PKUSIEMENS: t = −4.48, p < 0.001, df = 40), but recovered to baseline or even higher than baseline after distraction (IPCAS: t = 2.76, p = 0.0088, df = 40; PKUGE: t = 1.74, p = 0.09, df = 40; PKUSIEMENS: t = 2.43, p = 0.02, df = 40).

**Figure 1.**
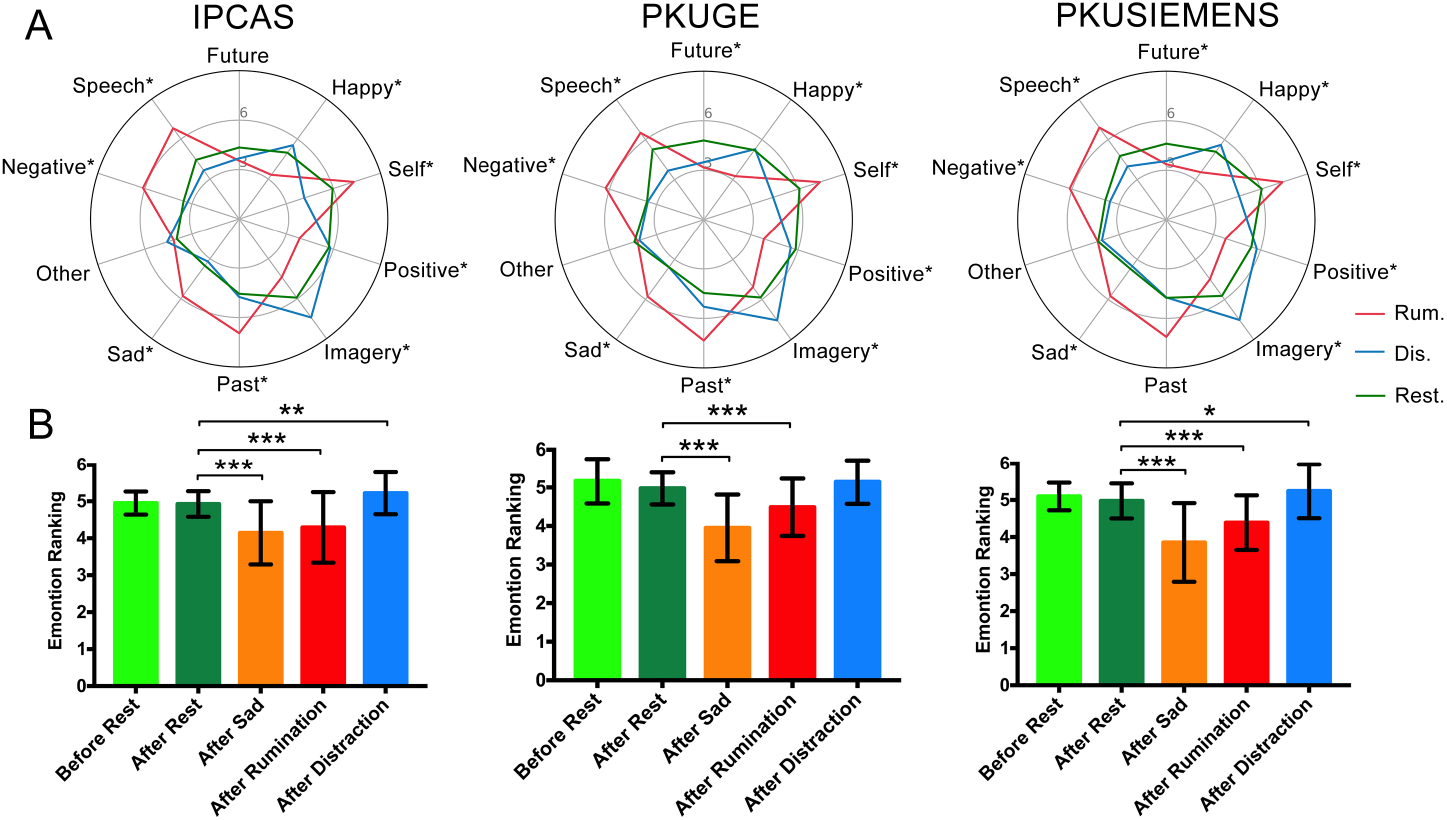
Self report behavior results. A) the form and content of thinking during rumination, distraction and resting states. Each axis stands for one dimension. *: significantly different among 3 states, p < 0.05, Bonferoni corrected. B) the emotional ratings (happy vs. unhappy) before resting state and after each states. *: p < 0.05; **: p <0.01; ***: p < 0.001.

### DMN Functional Connectivities

After the negative autobiographical memory induction, participants were either induced into rumination or distraction states. We first compared overall FC strengths within the DMN between rumination and distraction states. Results revealed overall decreased FC within the DMN during rumination as compared to distraction. This pattern was replicated on all 3 scanners (IPCAS: t = −2.27, p = 0.028, df = 39; PKUGE: t = −2.25, p = 0.030, df = 39; PKUSIEMENS: t = −3.33, p = 0.002, df = 39, see Figure 2).

**Figure 2.**
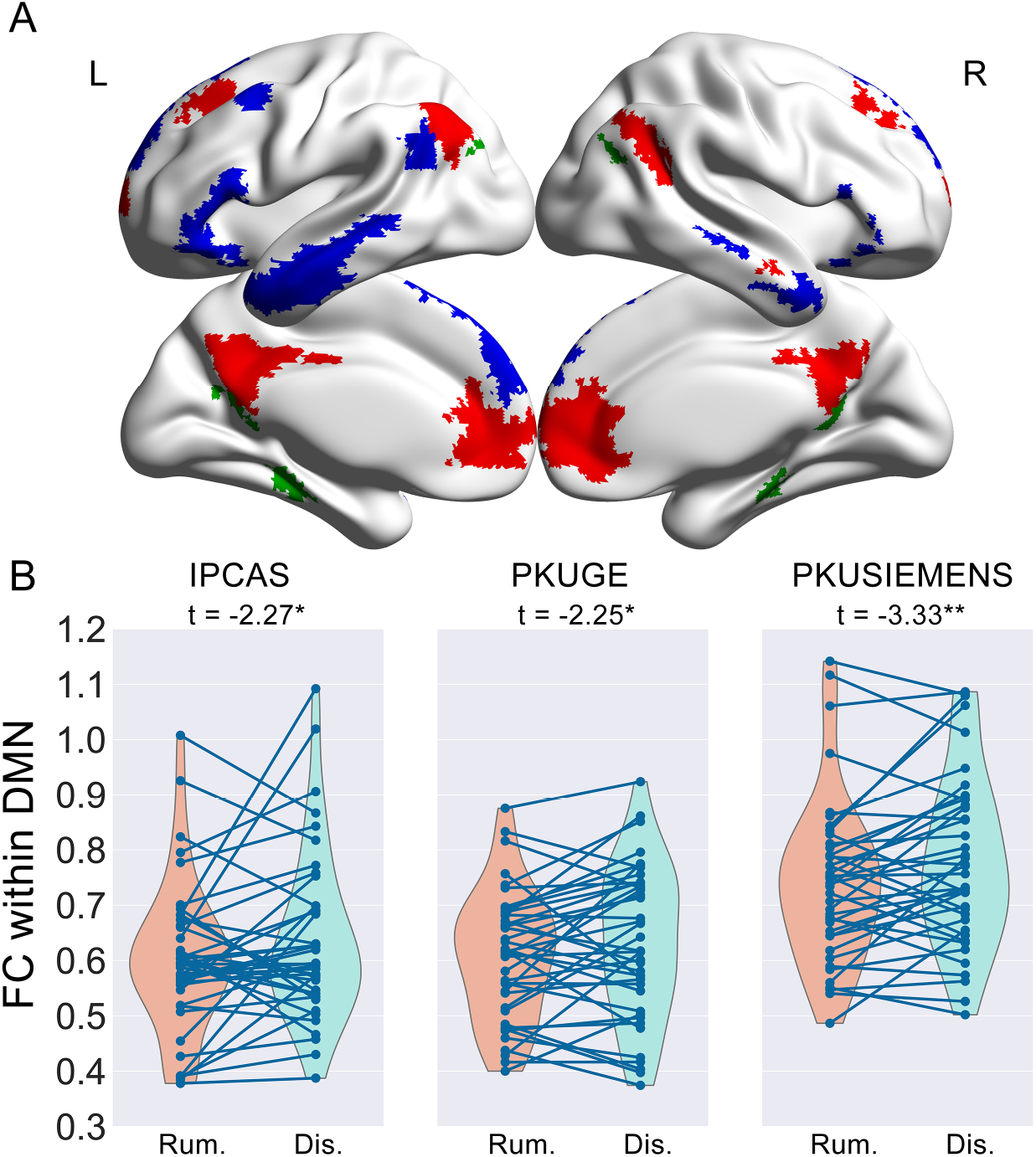
The differences of DMN FCs between rumination and distraction state across all 3 scanners. FCs have been adjusted for head motions. A) the DMN atlas used in the present study. Red: core subsystem; Blue: DMPFC subsystem; Green: MTL subsystem. L: left hemisphere; R: right hemisphere. B) Overall FC differences within DMN between rumination and distraction. *: p <0.05; **: p < 0.01. Rum.: rumination state; Dis.: distraction state.

### FCs Within and Between Subsystems of DMN

We further compared FCs within and between 3 subsystems of DMN in rumination and distraction states. Results revealed a reproducible but dissociate pattern. Specifically, FC between core and MTL subsystems was significantly enhanced in the rumination state as compared to distraction state (IPCAS: t = 1.30, p = 0.20, df = 39; PKUGE: t = 2.36, p = 0.023, df = 39; PKUSIEMENS: t = 2.88, p = 0.006, df = 39). FC between core and DMPFC subsystem were significantly decreased in rumination state as compared to distraction state (IPCAS: t = −3.02, p = 0.004, df = 39; PKUGE: t = −4.74, p < 0.001, df = 39; PKUSIEMENS: t = −3.37, p = 0.002, df = 39). FC within MTL subsystem in the rumination state was decreased versus the distraction state (IPCAS: t = −2.86, p = 0.007, df = 39; PKUGE: t = −1.67, p = 0.101, df = 39; PKUSIEMENS: t = −3.57, p = 0.001, df = 39, see Figure 3A and 3B). Further analysis showed that the system level findings were also remarkably reproducible at the ROI level. Some intriguing alterations in FCs between ROI pairs were revealed. FCs among left MTL, left ventral posterior inferior parietal lobule (vpIPL) and right PCC as well as right rostral medial prefrontal cortex (RMPFC) were elevated in rumination compared to the distraction state (Figure 4 & S3). FCs between left DMPFC and several ROIs of the core subsystem and FC between left and right MTL were decreased during rumination as compared to distraction states (FDR corrected p < 0.05, Figure S3). We also analyzed the whole brain FCs of left and right MTL and results revealed that left MTL was functional connected to more brain regions and more strongly during rumination as compared to distraction state (Figure S4).

**Figure 3.**
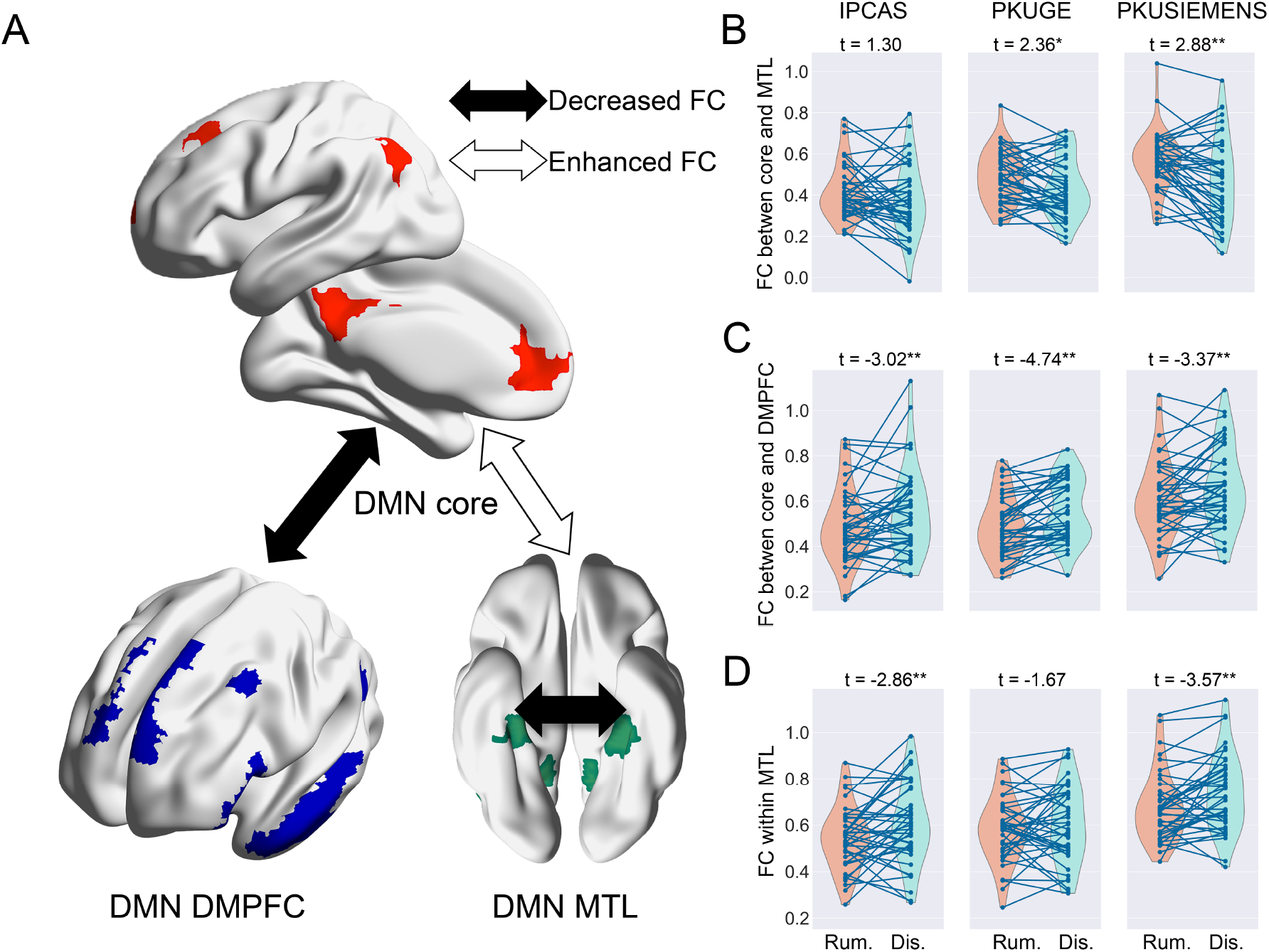
Differences of FCs among DMN subsystems between rumination and distraction state. A) Illustration of FC differences among DMN subsystems. FCs have been adjusted for head motions. B) An enhanced FCs between core and MTL subsystem in rumination state as compared to distraction state. C) Decreased FC between core and DMPFC subsystem in rumination state as compared to the distraction state. D) FC within MTL subsystem of DMN is decreased in rumination state as compared to distraction state. Note the reproducibility through all 3 scanners of these findings. **: significant after Bonferroni correction; *: significant uncorrected.

**Figure 4.**
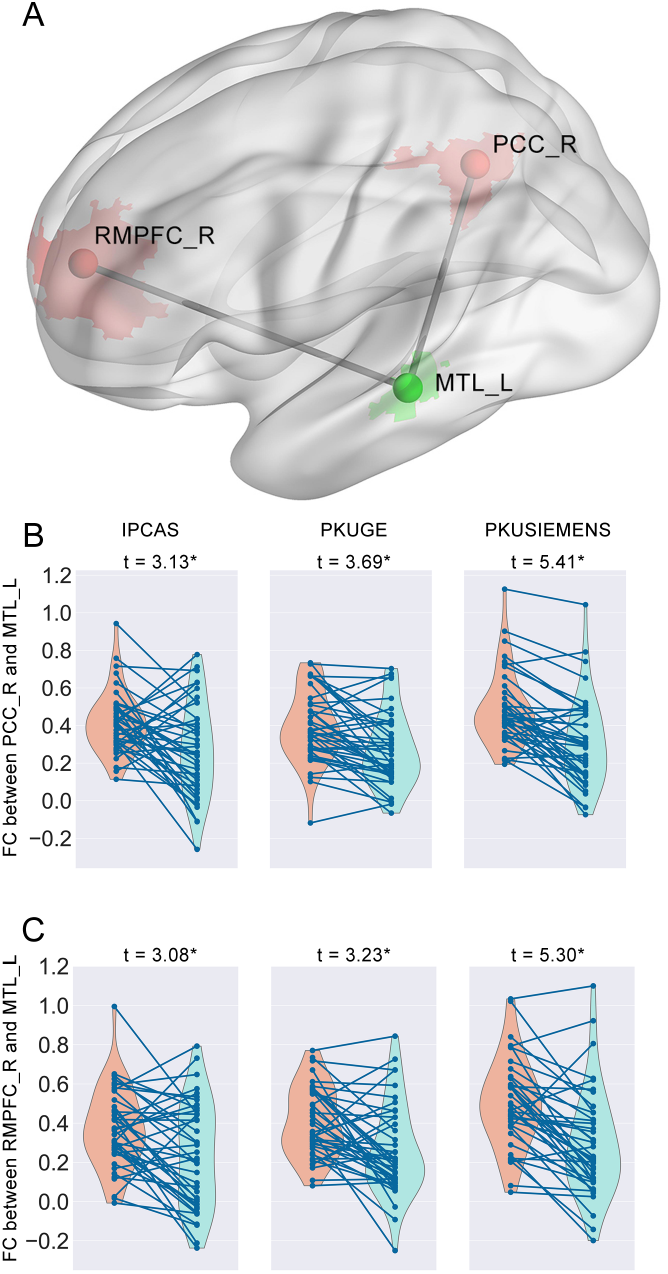
FCs between right PCC and left MTL as well as right RMPFC and left MTL revealed reproducible differences between rumination and distraction states. FCs have been adjusted for head motions. A) The ROIs we used to extract brain activity time courses and ROI pairs showing significant differences. B) Violin plots depicting FCs’ differences on right PCC – left MTL pairs between rumination and distraction states across 3 scanners. C) Violin plots depicting FCs’ differences on right RMPFC – left MTL pairs between rumination and distraction states across 3 scanners. Abbreviations: PCC: posterior cingulate cortex; MTL: medial temporal lobe; RMPFC: rostral medial prefrontal cortex; ROI: region of interest; FC: functional connectivity.

### ISCs within DMN

The comparison of ISCs within DMN revealed a decreased ISC within MTL subsystem in rumination as compared to distraction. This effect was significant on the PKUGE scanner (t = −3.01, p = 0.005, df = 40) whereas a similar, albeit non-significant, pattern was detected on the others (IPCAS: t = −1.55, p =0.129, df = 40; PKUSIEMENS: t = −1.41, p = 0.168, df = 40, see Figure 5). Further ROI level analysis found that both hemispheres’ retrosplenial cortexes (RSC)’ ISCs were significantly decreased and other ROIs in the MTL subsystem showed a similar pattern except for the right MTL (Figure S5).

**Figure 5.**
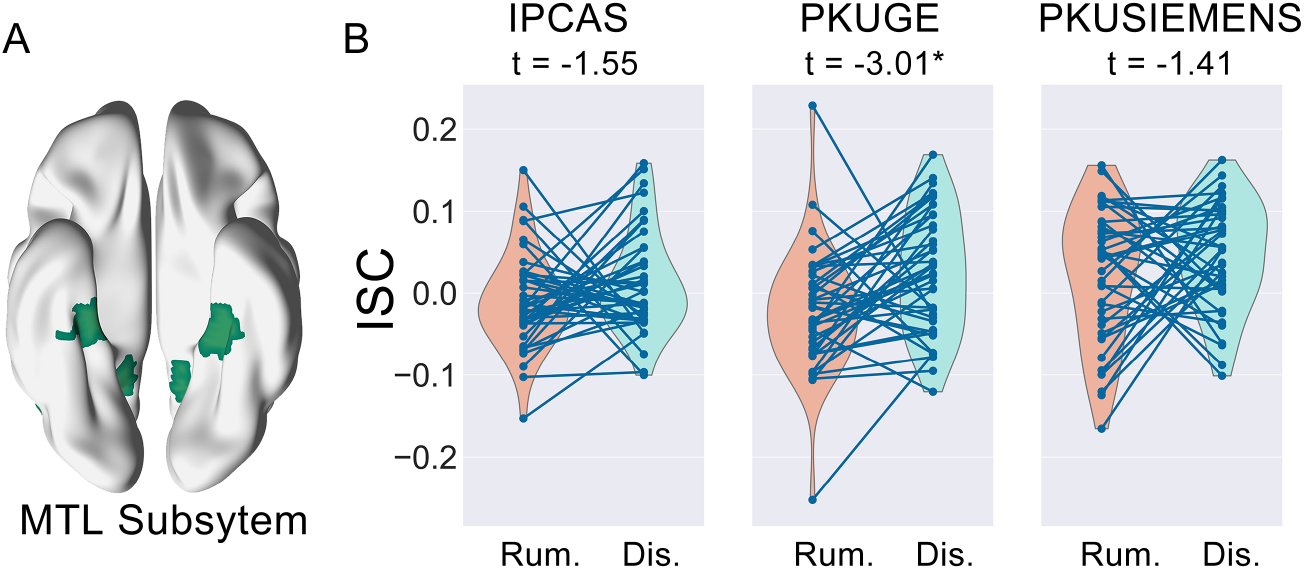
Differences of Inter-subject correlation (ISC) in the MTL subsystem between rumination and distraction states. A) illustration of the MTL subsystem ROIs. B) Decreased ISC within the MTL subsystem. *: statistically significant.

## Discussion

Here we examined DMN FC and ISC patterns during rumination state with a continuous mental state paradigm. We found overall decreased FCs within DMN during rumination as compared to distraction state. Further DMN subsystem analysis revealed a more nuanced pattern: compared to distraction state, FC between the core and MTL subsystems was enhanced whereas FC between the core and the DMPFC subsystems as well as FC within the MTL subsystem was decreased. Furthermore, ISC within the MTL subsystem was also decreased. Of note, our results were robustly reproducible across 3 different scanners.

Participants’ subjective reports revealed striking reproducibility across all 3 scanners. Emotional ratings showed that participants were in a dysphoric mood after recalling negative autobiographical events, indicating a successful negative affect induction. Because rumination is typically accompanied by psychological distress (Nolen-Hoeksema S *et al*. 2008), we believe this induction functions as an effective facilitation. Furthermore, reported forms and contents of thinking were generally in line with rumination’s phenomenological features: self-focused, negative and past oriented (Nolen-Hoeksema S *et al*. 2008; Andrews-Hanna J *et al*. 2013; Christoff K *et al*. 2016). To sum, behavioral self-ratings provided support for our paradigm in successfully inducing participants into a rumination state.

Our finding that FCs within DMN were decreased during rumination may seem counter-intuitive, but finer subsystem level analysis indicates this decrease may be driven by a dissociated FC pattern among different subsystems. Task based studies on rumination convergently found increased activity in the core subsystem as compared to distraction, but evidence regarding the other two subsystems has been less consistent (Zhou HX et al. 2019). Previous rumination state studies indicated FCs between core subsystem region (PCC) and rest DMN regions may best differentiate rumination state and resting state (Berman MG *et al*. 2014). Resting state research mainly implicated scale-measured rumination trait with FCs of core subsystem region (RMPFC) (Greicius MD *et al*. 2007; Zhu X *et al*. 2012; Kaiser RH *et al*. 2016). Whereas these previous studies provided informative regional evidence, our results provided an overall network-wise interaction pattern. Critically, our primary finding adds to the recent challenge to the prevailing notion that rumination causes the abnormally enhanced within DMN-FCs in MDD patients (Yan CG et al. 2019), indicating rumination’s DMN neural underpinnings are more nuanced than previous hypothesis have predicted.

One particularly intriguing finding was the enhanced FC between the core subsystem and the MTL subsystem during rumination as compared to distraction state. This is in line with the notion that rumination may be caused by the too strong influence the core subsystem exerts on the MTL subsystem (Bar M 2009; Christoff K *et al*. 2016). The core subsystem has been demonstrated to play an important role in processes such as self-referential perception, generation of affective meaning, and allowing individuals to construct and simulate alternative scenarios (Andrews-Hanna JR 2012; Roy M et al. 2012). On the other hand, the MTL subsystem was proposed to engage in the retrieval of autobiographical memory and the generation of novel “blocks” of spontaneous thoughts (Bar M et al. 2007; Bar M 2009). Further ROI level analysis showed this effect was primarily driven by FCs among RMPFC, PCC and left MTL, which were all key regions underlying above-mentioned psychological processes (Figure 4). RMPFC and both MTLs were also demonstrated to be especially relevant to affect processes (Fox KCR *et al*. 2018). Our results implicated that the over-constraint from core DMN regions (RMPFC and PCC) to the MTL subsystem regions (MTL) may lead to the key phenomenological features of rumination: repetitive, self-focus and negative valence (Christoff K *et al*. 2016; Marchetti I et al. 2016; Fox KCR *et al*. 2018).

We also found decreased FC between the core and DMPFC subsystems. The DMPFC subsystem was proposed to be linked to the theory of mind process, mental simulations and present-oriented thoughts (Buckner RL and DC Carroll 2007; Andrews-Hanna JR *et al*. 2014), but less involved in the generation of spontaneous thoughts, affective processes and past or future oriented thoughts (Andrews-Hanna JR, JS Reidler, J Sepulcre*, et al*. 2010; Christoff K *et al*. 2016). Distraction is a present oriented, unrelated-to-self and non-affective mental state and has little to do with autobiographical processes (Morrow J and S Nolen-Hoeksema 1990). These features may lead to a stronger connection between the core and DMPFC subsystems during distraction as compared to rumination. Further ROI level analysis showed that the subsystem level difference was primarily driven by decreased FC between DMPFC and brain regions of the core subsystem. DMPFC projects less to the so-called “limbic system” and is associated with symbolic and universal rather than affective and motivational representations, which may be more involved in the distraction state (Ongur D and JL Price 2000; Olsson A and KN Ochsner 2008). Thus, it is possible that during rumination, brain regions of the DMPFC subsystem, especially DMPFC, cooperate less with the core subsystem because rumination consists of less symbolic, non-emotional and present oriented representations as compared to distraction. Together, these findings implicate the specific interaction modes among DMN subsystems that may characterize this very form of spontaneous thought: rumination.

We found that FCs within the MTL subsystem were decreased when participants were ruminating versus when they were distracted. This may be represented by the lateralization of both MTLs. Although they were both considered to be essential in autobiographical memory retrieving and generating spontaneous thoughts (Bar M 2009; Rugg MD and KL Vilberg 2013), the left MTL was posited to be engaged in processing of verbal materials more than non-verbal materials (Martin A 1999; Golby AJ et al. 2002). Rumination is primarily in a verbal rather than imagery form, so left MTL may activate differently from right MTL during rumination, which probably reduces their FC. Indeed, our results showed left MTL was more widely and densely functionally connected to core DMN regions than right MTL during rumination as compared to distraction, indicating its more critical role in rumination (Figure S4).

Additionally, we found reduced ISCs within the MTL subsystem. Further ROI level analysis indicated an overall ISC decrease except for right MTL. Recent evidence revealed that similar DMN responses could also be observed when individuals were performing similar internal mental processes such as memory retrieval narrative understanding (Chen J *et al*. 2017; Nguyen M et al. 2019). It is possible that similar external stimuli or internal thinking contents elicit shared DMN patterns. Compared to distraction, contents of thought are more individualized during rumination. Each individual has unique concerns or understanding of themselves or of negative events, and these thoughts depend on the novel materials generated by the MTL subsystem, which may be reflected by the lower consistency of activity across participants.

Reproducibility is a current major concern in functional MRI (Poldrack RA *et al*. 2017). In the neuroimaging field, a fundamental notion for reproducibility would be measuring same subjects with different scanners and test to what extent results can be generalized(Nichols TE et al. 2017). Despite using multiple scanners and sites, our results still yielded a remarkable reproducibility. This consistency indicates that the effects we report are likely prominent and stable. Furthermore, raw imaging data and codes were made openly available to allow independent replication of our findings, as enhancements help improve the transparency and reproducibility of our present study(Gorgolewski KJ and RA Poldrack 2016). Because rumination has long been deemed as an important phenomenological feature and risk factor for MDD (Nolen-Hoeksema S *et al*. 2008), we wish these highly reproducible findings may act as future neuroimaging biomarkers of this common and disabling psychiatric condition.

It should be noted that our results were exclusively based on a young healthy adult sample and cannot be generalized to clinical MDD patients. Future studies should be conducted with clinical samples to obtain a full understanding of the pathological role of rumination in MDD.

In summary, with repeated measures, we discerned more nuanced and dissociated DMN network underpinnings of rumination. Our current results offer direct evidence on how DMN regions synchronize within and among subjects to underlie rumination, thus shedding new light on understanding the complex relationship between rumination and DMN abnormalities in MDD patients.

## Acknowledgments

This work was supported by the National Key R&D Program of China (grant number: 2017YFC1309902 to CY), the National Natural Science Foundation of China (grant number: 81671774, 81630031 to CY), the Hundred Talents Program and the 13th Five-year Informatization Plan of Chinese Academy of Sciences (grant number: XXH13505 to CY), Beijing Municipal Science & Technology Commission (grant number: Z161100000216152 to CY). The authors appreciate the editorial assistance and support of Dr. Francisco X. Castellanos and would like to thank Dr. Men Weiwei for his technical supports during data collection.

