## Supporting Information for "The Subsystem Mechanism of Default Mode Network Underlying Rumination: a Reproducible Neuroimaging Study"

**Chen et al.**

Table S1. The stimuli used to instruct rumination and distraction states.

| Instructions | Condition |
|---|---|
| Think: Analyze your personality to understand why you feel so depressed in the events you just memorized | Rumination |
| Think: The kind of person you are revealed by the events you just memorized. How similar/different you are relative to other people？ | Rumination |
| Think: Why do I encounter these events other people don’t？ | Rumination |
| Think: Why can’t I handle things better in these events I just memorized？ | Rumination |
| Think about: The layout of a typical classroom | Distraction |
| Think about: Raindrops sliding down a windowpane | Distraction |
| Think about: Clouds forming in the sky | Distraction |
| Think about: A train stopped at a station | Distractions |

Table S2. Scales participants used to report the form and contents of their thinking during different mental states. Participants reported with a 9-point Likert scale (1: completely absent; 9: almost all).

| Serial number | Scale items |
|---|---|
| 1 | I was thinking about the PAST |
| 2 | I was thinking about the FUTURE |
| 3 | I was thinking about MYSELF |
| 4 | I was thinking about OTHERS |
| 5 | What I thought were POSITIVE |
| 6 | What I thought were NEGATIVE |
| 7 | What I thought were in the form of IMAGE |
| 8 | What I thought were in the form of SPEECH |
| 9 | What I thought made me feel HAPPY |
| 10 | What I thought made me feel SAD |

Table S3. ROIs used in the present study.

| Serial Number | Abbreviation | ROI | Hemisphere | subsystem |
|---|---|---|---|---|
| 1 | pIPL_L | posterior inferior parietal lobule | Left | core |
| 2 | SFS_L | superior frontal sulcus | Left | core |
| 3 | PCC_L | posterior cingulate cortex | Left | core |
| 4 | RMPFC_L | rostromedial prefrontal cortex | Left | core |
| 5 | rSTS_R | right rostral superior temporal sulcus | Right | core |
| 6 | pIPL_R | posterior inferior parietal lobule | Right | core |
| 7 | SFS_R | superior frontal sulcus | Right | core |
| 8 | PCC_R | posterior cingulate cortex | Right | core |
| 9 | RMPFC_R | rostromedial prefrontal cortex | Right | core |
| 10 | LTC_TP_L | temporal pole of lateral temporal cortex | Left | DMPFC |
| 11 | TPJ_L | left temporoparietal junction | Left | DMPFC |
| 12 | DMPFC_L | dorsomedial prefrontal cortex | Left | DMPFC |
| 13 | pDLPFC_L | left posterior dorsolateral prefrontal cortex | Left | DMPFC |
| 14 | IFG_L | inferior frontal gyrus | Left | DMPFC |
| 15 | LTC_R | lateral temporal cortex | Right | DMPFC |
| 16 | TP_R | temporopolar cortex | Right | DMPFC |
| 17 | DMPFC_R | dorsomedial prefrontal cortex | Right | DMPFC |
| 18 | IFG_R | inferior frontal gyrus | Right | DMPFC |
| 19 | vpIPL_L | ventral posterior inferior parietal lobule | Left | MTL |
| 20 | RSC_L | retrosplenial cortex/ventral posterior cingulate | Left | MTL |
| 21 | MTL_L | medial temporal lobe | Left | MTL |
| 22 | vpIPL_R | ventral posterior inferior parietal lobule | Right | MTL |
| 23 | RSC_R | retrosplenial cortex/ventral posterior cingulate | Right | MTL |
| 24 | MTL_R | medial temporal lobe | Right | MTL |

Table S4. Subjective ratings on all dimensions of thinking contents during rumination and distraction state. Means and standard deviations as well as t and p stats were listed.

| | IPCAS | | | | PKUGE | | | | PKUSIEMENS | | | |
|---|---|---|---|---|---|---|---|---|---|---|---|---|
| | Rumination (mean ± SD) | Distraction (mean ± SD) | t | p | Rumination (mean ± SD) | Distraction (mean ± SD) | t | p | Rumination (mean ± SD) | Distraction (mean ± SD) | t | p |
| Past | 6.95±1.50 | 4.76±2.06 | 5.530 | <0.001 | 7.34±1.13 | 5.29±2.14 | 5.313 | < 0.001 | 7.12±1.36 | 4.73±2.05 | 6.716 | < 0.001 |
| Future | 3.54±1.75 | 3.68±1.57 | -0.458 | 0.649 | 3.17±1.58 | 3.49±1.72 | -0.880 | 0.384 | 3.37±1.58 | 3.56±1.52 | -0.628 | 0.534 |
| Self | 7.29±1.23 | 4.15±2.08 | 7.915 | <0.001 | 7.39±0.95 | 4.41±2.10 | 7.895 | < 0.001 | 7.39±0.89 | 4.63±1.92 | 8.376 | < 0.001 |
| Other | 4.17±1.96 | 4.59±2.04 | -1.089 | 0.283 | 4.22±1.72 | 4.10±1.84 | 0.299 | 0.766 | 4.37±2.02 | 4.10±1.66 | 0.647 | 0.521 |
| Positive | 3.85±1.68 | 5.80±1.40 | -6.497 | <0.001 | 3.83±1.63 | 5.54±1.27 | -5.713 | < 0.001 | 3.78±1.52 | 5.76±0.99 | -7.735 | < 0.001 |
| Negative | 6.12±1.78 | 3.29±1.33 | 8.898 | <0.001 | 6.24±1.64 | 3.56±1.27 | 7.182 | < 0.001 | 6.15±1.46 | 3.59±1.36 | 8.409 | < 0.001 |
| Image | 4.39±2.22 | 7.41±1.36 | -7.611 | <0.001 | 5.07±2.31 | 7.56±1.21 | -5.853 | < 0.001 | 4.49±2.06 | 7.54±1.38 | -8.733 | < 0.001 |
| Image | 6.80±1.93 | 3.66±2.09 | 7.682 | <0.001 | 6.51±1.83 | 3.68±2.32 | 7.028 | < 0.001 | 6.90±1.80 | 4.00±2.05 | 8.197 | < 0.001 |
| Speech | 3.32±1.35 | 5.54±1.12 | -8.455 | <0.001 | 3.27±1.16 | 5.29±1.17 | -8.595 | < 0.001 | 3.56±1.29 | 5.61±1.18 | -8.140 | < 0.001 |
| Happy | 5.80±1.65 | 3.22±1.27 | 8.914 | <0.001 | 5.78±1.54 | 3.59±1.36 | 7.448 | < 0.001 | 5.76±1.48 | 3.51±1.36 | 8.175 | < 0.001 |
| Sad | 6.95±1.50 | 4.76±2.06 | 5.530 | <0.001 | 7.34±1.13 | 5.29±2.14 | 5.313 | < 0.001 | 7.12±1.36 | 4.73±2.05 | 6.716 | < 0.001 |

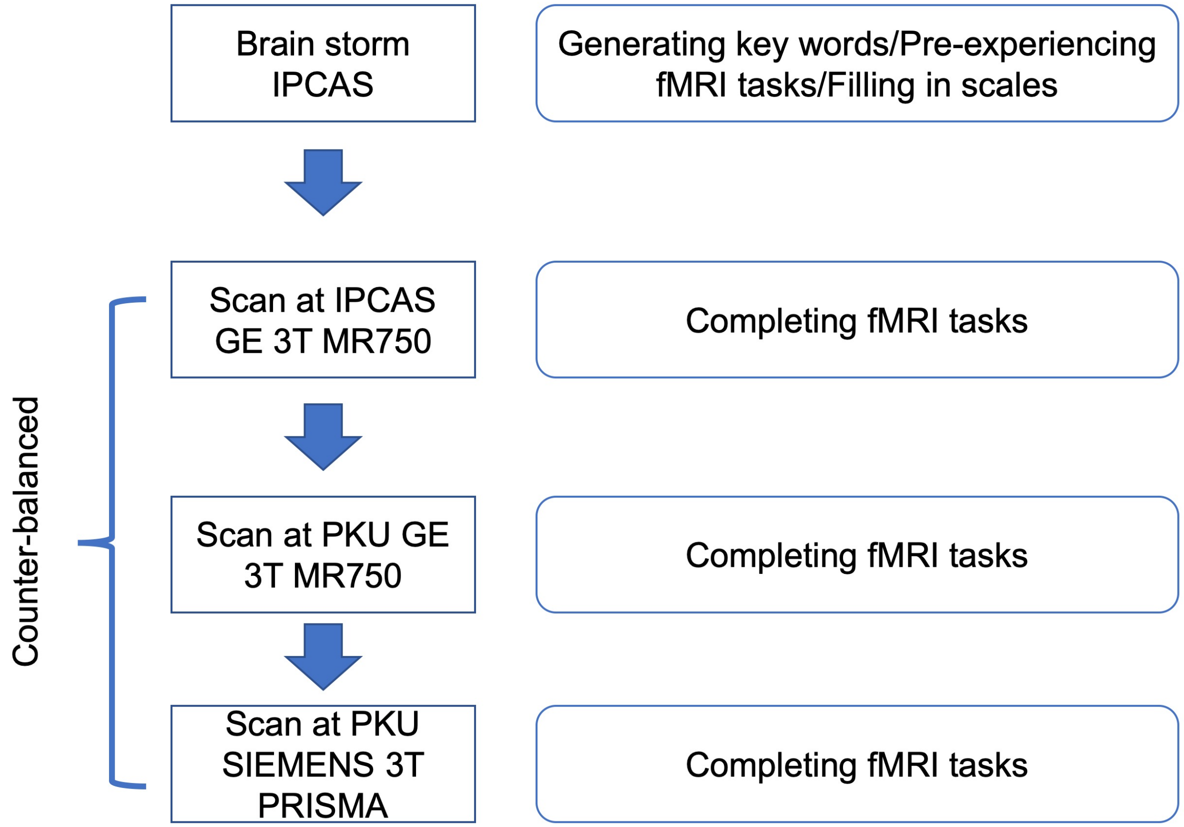


Figure S1. Sequence and tasks of brain storm and three scans on different scanners. The identical fMRI task was repeated for 3 times on 3 different scanners and the order was counter-balanced.

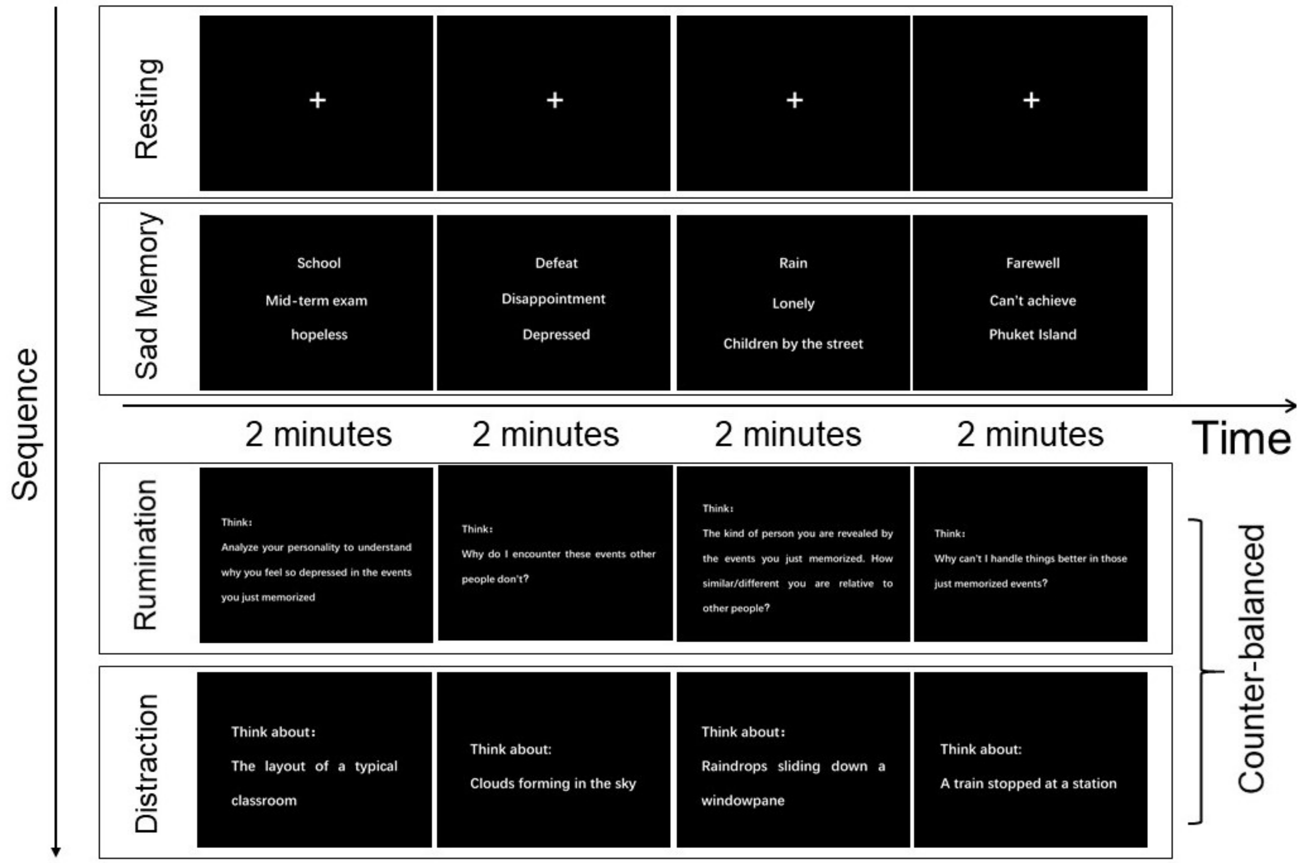


Figure S2. Flow chart and illustration of fMRI task. The fMRI scan consists of 4 runs. The first run was an 8-minute resting state, and a "+" symbol will be displayed on the screen. After that, participants underwent a negative autobiographical memory session (sad memory). Four sets of key words which were generated during brain storm were displayed on the screen sequentially and the order of them were randomized across participants. This session functioned as a mood induction. After this sad memory session, participants were induced either to rumination or distract. During rumination, questions modified from the Rumination Induction Task (RIT) were displayed sequentially on the screen. Four objective scenes were displayed in the distraction state. The 3 mental states (sad memory, rumination, distraction) consists of 4 stimuli which lasted for 2 minutes and no inter stimuli intervals. The order of these stimuli was randomized across participants. The details of the instructions we used in rumination and distraction states can be found in Table S3.

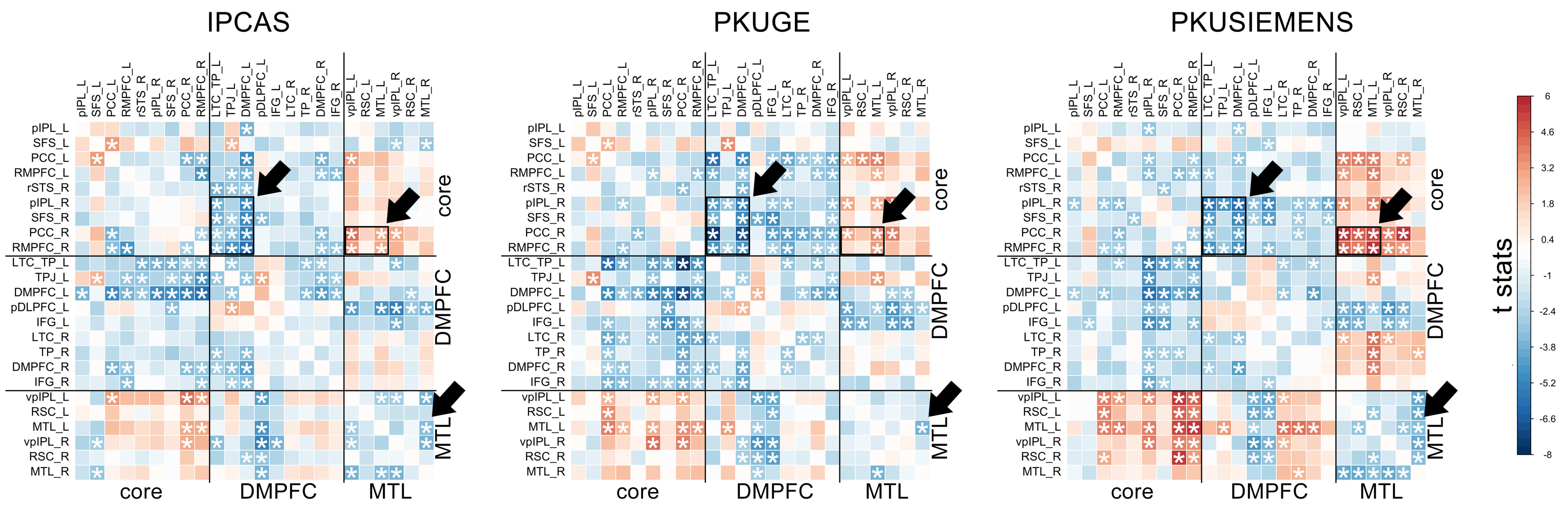


Figure S3. Heatmap showing t stats of FC differences between rumination and distraction states on the ROI level. Note the overall reproducible pattern across 3 scanners. FCs between left dorsal prefrontal cortex (DMPFC) and temporal pole of left lateral temporal cortex (LTC_TP) and several regions of interest (ROI) of the core subsystem, FCs among left ventral posterior inferior parietal lobe (vpIPL), left medial temporal lobe (MTL) and right posterior cingulate cortex (PCC), FC between left and right MTLs revealed a reproducible altered pattern in rumination as compared to distraction. All ROIs and their corresponding abbreviation please refer to Table S3. *: FDR corrected $p < 0.05$.

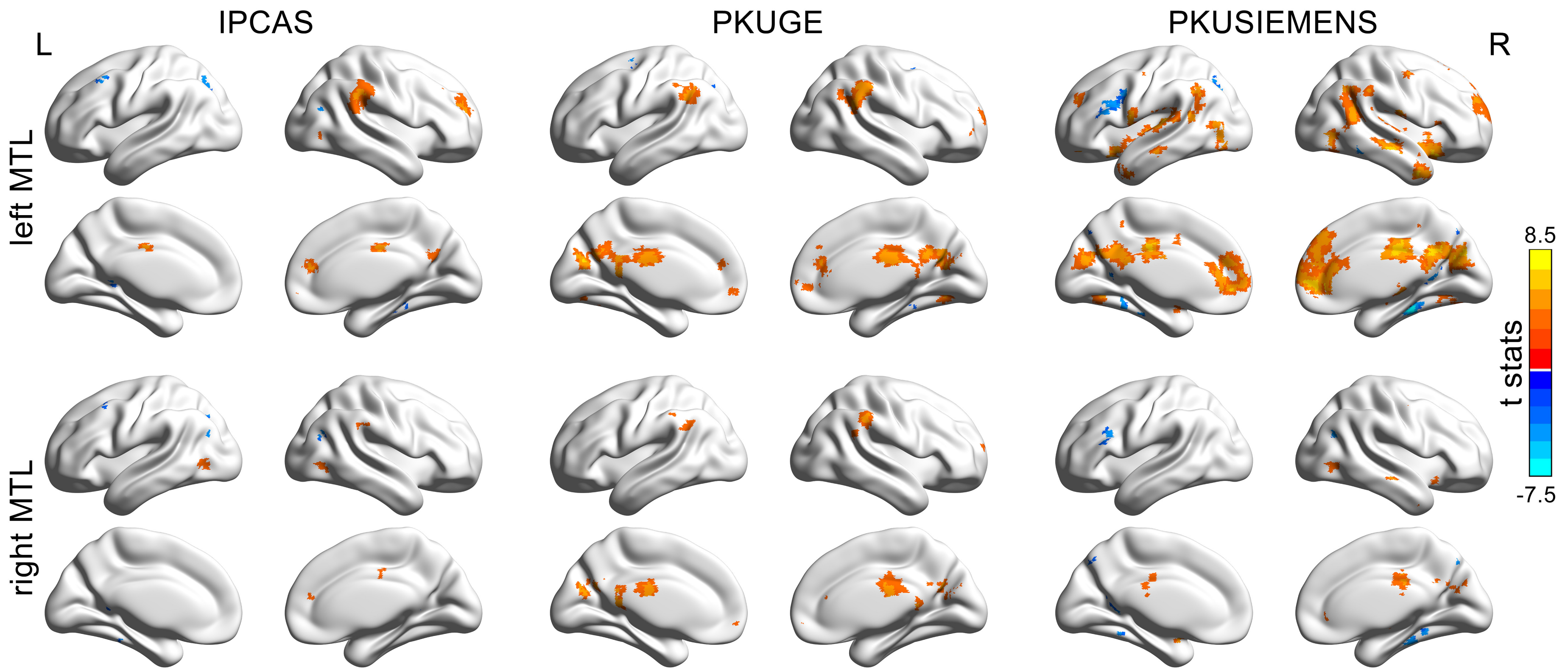


Figure S4. FC difference maps of left and right medial temporal lobe (MTL) between rumination and distraction states across whole brain ($p < 0.001$, uncorrected). Note the much wider elevated functional connectivity (FC) patterns of left MTL during rumination as compared to the right MTL. This indicates the more lateralized DMN FC pattern of both MTLs during rumination.

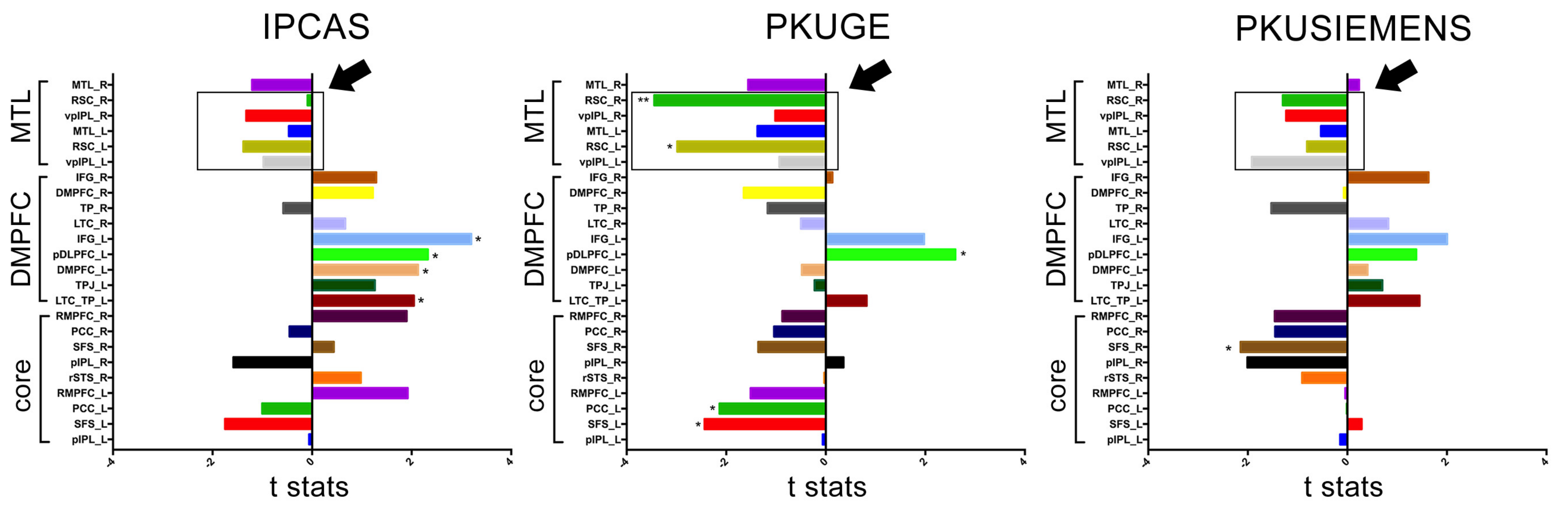


Figure S5. Bar plot depicting t stats of inter-subject correlation (ISC) differences of each regions of interest (ROI) from all 3 subsystems between rumination and distraction states. ISCs of both retrosplenial cortexes (RSC) were significantly decreased during rumination as compared to distraction state. Other ROIs within the medial temporal lobe (MTL) subsystem revealed a similar pattern except for right MTL. *: significant but not survive the FDR correction; **: significant after FDR correction.
